# ADAR2-repressed RNA editing: a novel mechanism contributing to t (8:21) AML leukemogenesis

**DOI:** 10.1101/2021.10.18.464918

**Authors:** Mingrui Guo, Tim Hon Man Chan, Omer An, Yangyang Song, Zi Hui Tan, Vanessa Hui En Ng, Xinang Cao, Wee Joo Chng, Motomi Osato, Henry Yang, Qiling Zhou, Leilei Chen, Daniel G. Tenen

**Affiliations:** Cancer Science Institute of Singapore, National University of Singapore, 117599 Singapore, Singapore; YLL School of Medicine, National University of Singapore, 119228 Singapore, Singapore; Department of Laboratory Medicine, Molecular Diagnosis Centre, National University Health System, Singapore 119074, Singapore; Duke-NUS Medical School, Singapore 169857, Singapore; Department of Haematology-Oncology, National University Cancer Institute of Singapore, National University Health System, Singapore, Singapore; Department of Anatomy, Yong Loo Lin School of Medicine, National University of Singapore, Singapore 117594, Singapore; Harvard Stem Cell Institute, Harvard Medical School, Boston, MA 02115, USA

**Keywords:** AML, Leukemogenesis, RUNX, ADAR, A-to-I RNA editing, COG3, COPA

## Abstract

In the past decade, adenosine to inosine (A-to-I) RNA editing, which is catalyzed by adenosine deaminases acting on RNA (ADAR) family of enzymes ADAR1 and ADAR2, has been shown to contribute to the development and progression of multiple cancers; however, very little is known about its role in acute myeloid leukemia (AML) - the second most common type of leukemia making up 31% of all adult leukemia cases. Here, we found that ADAR2, but not ADAR1 and ADAR3, is specifically downregulated in core binding factor (CBF) AML with t(8;21) or inv(16). In t(8;21) AML, RUNX1-driven transcription of ADAR2 transcripts was found to be repressed by the RUNX1-ETO fusion protein. Forced overexpression of two ADAR2-regulated RNA editing targets COPA and COG3 indeed inhibits clonogenic growth of human t(8;21) AML cells. Further *in vivo* animal studies confirmed that ADAR2 could suppress leukemogenesis of t(8;21) AML through its RNA binding and editing capabilities. Our results suggest a novel RNA editing-mediated mechanism leading to t(8,12) AML.

**Key points:**

- ADAR2, but not ADAR1 and ADAR3, was specifically downregulated in CBF-AML
- RUNX1-ETO suppresses ADAR2 transcription in t(8;21) AML through binding on its promoter
- RNA editing capability of ADAR2 is essential for its repression of leukemogenesis in an AE9a mouse model

## Introduction

High throughput technologies such as RNA sequencing have revolutionized our understanding of global transcriptomic changes. RNA processing steps including alternative splicing, alternative polyadenylation, and RNA editing/modifications, significantly contribute to the composition and complexity of the transcriptome. Adenosine-to-inosine (A-to-I) RNA editing is the most prevalent type of RNA editing in mammals and is catalysed by the adenosine deaminase acting on RNA (ADAR) family of enzymes that recognise double-stranded RNAs (dsRNA)^1^. In vertebrates, the ADAR family consists of 3 members, ADAR1, ADAR2, and ADAR3^2^. ADAR1 and ADAR2 mediate the editing reaction which contributes to multilevel regulation of gene expression and activity, whereas ADAR3 has no documented deaminase activity^3^. A-to-I RNA editing not only alters the RNA sequence itself but also affect the cellular fate of RNA molecules. In principle, A-to-I editing sites can be found in both coding and non-coding regions. However, the vast majority of A-to-I editing sites are in introns and untranslated regions harbouring long and perfect dsRNA structures formed by inverted *Alu* repetitive elements^4^. Over the past decades, accumulating evidence suggests the dysregulated A-to-I editing as one of the key drivers for various cancers, particularly solid tumors^5-10^. In coding regions, RNA editing can lead to amino acid codon change. Differential editing of these protein-recoding sites are found to impact on human diseases, such as neurological diseases and cancer^6,8,10-12^. In cancer, only a few aberrant protein-recoding targets have been reported thus far, and they contribute to tumorigenesis largely through enhancing cancer promoting activity or repressing tumour suppressive activity^7,8,10,13^.

Till date, very limited efforts have been placed on understanding the role of RNA editing in haematological malignancies including acute myeloid leukemia (AML) which is the most common haematological malignancy characterized by the abnormal proliferation and differentiation of myeloid progenitor cells with high incidence and recurrence rates. Although there are several studies on ADAR1 and its mediated RNA editing events in multiple myeloma (MM)^14,15^ and chronic myeloid leukaemia (CML)^16-18^, the role of ADAR2 in AML and other haematological malignancies remain unknown. In this study, we analysed the expression profiles of three ADAR enzymes in AML patients with distinct molecular subtypes, from publicly available cDNA microarray^19^ and the TCGA RNA sequencing datasets^20^. Surprisingly, ADAR2, but not ADAR1 and ADAR3, was specifically downregulated in AML patients with t(8;21) or inv(16) and both of which belong to the core binding factor (CBF) AML comprising up to 12–15% of all AML cases^21-23^. CBF AML is characterized by the presence of either t(8;21)(q22;q22) or inv(16)(p13q22)/t(16;16), which results in the formation of *RUNX1-ETO* and *CBFβ-MYH11* fusion genes respectively. Core binding factors are transcription factors which are necessary in normal haematopoiesis and characterized by heterodimers of a DNA-binding unit CBFα (including three subunits RUNX1, RUNX2, RUNX3) and a non-DNA-binding unit CBFβ. Chromosomal translocations can alter DNA-binding capability and create alternate binding sites of the heterodimer, leading to disruption of normal transcription program and the consequent maturation arrest^24^. Unlike wildtype RUNX1, RUNX1-ETO fusion protein drive leukemogenesis through assembling transcriptional regulatory complexes, either as a repressor or an activator^25-29^. Although the presence of RUNX1-ETO (also named AML1-ETO) defines a precursor stage of leukaemia, additional molecular events are required for transformation^30,31^. In this study, we report that RUNX1-ETO fusion protein represses RUNX1-driven transcription of *ADAR2* transcripts. The functional investigation of ADAR2 and ADAR2-mediated RNA editing in human t(8;21) AML cells and a t(8;21) AML mouse model uncovered that ADAR2 could suppress leukemogenesis of t(8;21) AML through its RNA binding and editing capabilities. This is the first time that ADAR2 and its-mediated RNA editing events are linked to leukemogenesis of t(8;21) AML.

## Materials and methods

### Clinical tissue samples

Primary AML and matched normal knee samples were obtained from the CenTRAL (Molecular) Leukaemia Tissue Bank, with approvals from Institutional Review Board, National University of Singapore, and signed patient informed consent. In this study, “normal” samples refer to samples harvested from the knee samples of healthy individuals.

### Mice

C57BL/6J mice were obtained from The Jackson Laboratory. All mice were housed in a sterile barrier facility within the Comparative Medicine facility at the National University of Singapore under housing condition of 22 °C temperature, 50% humidity and a 12:12 light/dark cycle. All mice experiments performed in this study were approved by Institutional Animal Care and Use Committee of National University of Singapore. In this study, 8-to 16-week old male mice were used for bone marrow harvesting or transplantation; 8-to 16-week old female mice were used for time mating. See method detail for more information.

### Establishment of Kasumi-1 stable cell lines

To generate Kasumi-1 cells stably expressing *ADAR2, ADAR2* mutants, wildtype or edited forms of *COPA* and *COG3*, 4.5 µg VSV-G and 4.5 µg expression constructs were co-transfected into GP2-293 cells cultured in T75 flask for virus generation. Supernatant containing lentivirus was harvested and filtered using 0.45um syringe filter (Sartorius, cat.no. 16537) at 48 and 72 hours after transfection. These two portions of supernatant were mixed and aliquoted for virus transduction or stored in -80°C for subsequent usage. Virus transduction was performed using RetroNectin® Recombinant Human Fibronectin Fragment (Takara, cat.no. T100A/B) according to the manufacturer’s instructions (with centrifugation). At 72 hours after virus transduction, cells were selected using puromycin (2 µg/ml) for 72 hrs or by FACS sorting.

### Luciferase reporter assays

The firefly luciferase reporter gene in the pGL3 vector is driven by human *ADAR2* promoter regions containing different RUNX site portions. As an internal control plasmid for co-transfections, the pRL-null construct encoding a Renilla luciferase gene (Promega, cat.no. E2231) was used. Firefly and Renilla luciferase activities were determined 24 hours post transfection with the dual-luciferase reporter assay system (Promega, cat.no. E1910). Firefly luciferase readings were normalized against internal control Renilla luciferase and calculated as fold differences against the activity obtained from cells transfected with empty vector.

### Analysis of RNA editing by Sanger-sequencing

To amplify regions containing *COG3 I635V* or *COPA I164V* site, cDNA from different cells were used for PLATINUM GREEN HS PCR 2X Master PCR amplification (Thermo fisher scientific. cat.no. 13001014. Purified PCR products were sent for direct sequencing, and the result was visualized with SnapGene software (SnapGene®, San Diego, CA,USA). The frequencies of *COPA* and *COG3* editing were calculated based on the peak area of adenosine and guanosine determined by SnapGene. Sequence of primers are listed in Supplemental Table 2.

### RUNX1-ETO9a primary leukemia model

LSK population from fetal liver cells from embryos at E14.5– E16.5 stage was used for AE9a retrovirus transduction. At 7 days after transduction, fifty thousands of FACS-sorted GPF positive cells were transplanted into sub-lethally (6.5 Gy) γ-irradiated C57BL/6J mice through retro-orbital injection. Moribund mice were euthanized with CO_2_ and dissected for spleen, vertebrae, femur, tibia, and hip collection. A secondary transplantation was performed through transplanting two hundred thousand bone marrow cells from the first round of transplantation into sub-lethally (6.5 Gy) γ-irradiated C57BL/6J mice through retro-orbital injection. Moribund mice were euthanized with CO_2_ and dissected for spleen, vertebrae, femur, tibia, and hip collection. The bone marrow cells were stored in -80°C for subsequent usage.

### Rescue of *ADAR2*/*ADAR2* mutant in AE9a mouse model

*ADAR2* or *ADAR2* mutants retrovirus were transduced into BM cells from AE9a leukemic mice in the 2^nd^ transplantation. At day 2 after transduction, fifty thousands of cells were transplanted into sub-lethally (6.5 Gy) γ-irradiated C57BL/6J mice through retro-orbital injection for Peripheral blood (PB) collection and survival monitoring. PB samples (∼100 μl per mouse) were collected in EDTA-coated capillary tube (Drummond Scientific, cat.no. 1-000-800/12) by submandibular venipuncture with 5-mm Goldenrod animal lancets (Braintree Scientific, cat.no. GR5MM). Counts of nucleated cells was performed using a NIHOKODEN auto blood cell counter under Pre-dilute 20 µl mode. Moribund mice were euthanized with CO_2_ and dissected for spleen, vertebrae, femur, tibia, and hip collection. BM cells harvested from moribund mice were cytospun and stained with Giemsa’s azur-eosin-methylene blue solution (Merck, cat.no. 109204).

### Statistical analysis

The statistical significances were assessed by two-tailed Student’s *t*-test using the Excel unless otherwise specified.

Additional details are described in the supplemental information.

## Results

### *ADAR2* is significantly downregulated in CBF AML

To understand the role of ADARs and their mediated RNA editing in AML, we first utilized a publicly available microarray dataset^19^ which determined the gene-expression profiles in samples of peripheral blood or bone marrow from 285 patients with AML using Affymetrix U133A GeneChips. Intriguingly, although previous studies reporting the role of ADAR1 and ADAR1-regulated RNA editing events are implicated in haematological malignancies, we found that *ADAR2*, but not *ADAR1* and *ADAR3*, was significantly downregulated in CBF AML patients (**Figure 1A**). We next analysed the RNA sequencing (RNA-seq) data from TCGA. Despite a small same size, we found that the expression level of *ADAR2*, but not *ADAR1* and *ADAR3*, was significantly lower in t(8;21) or inv16 AML patients than the non-t(8;21) and inv16 AML patients (**Figure 1B, Supplemental Figure 1**). To experimentally validate our findings, we examined the expression level of *ADAR2* in an in-house AML cohort and healthy controls. In agreement with the above-mentioned expression analyses, *ADAR2* was found to be downregulated in t(8;21) AML patients when compared to CBF-negative patients and healthy controls (**Figure 1C**). Altogether, ADAR2 is most likely to be the only ADAR enzyme showing expression fluctuations among different AML subtypes and its selective downregulation in CBF AML patients suggests a previously undescribed mechanism which may lead to leukemogenesis.

**Figure 1.**
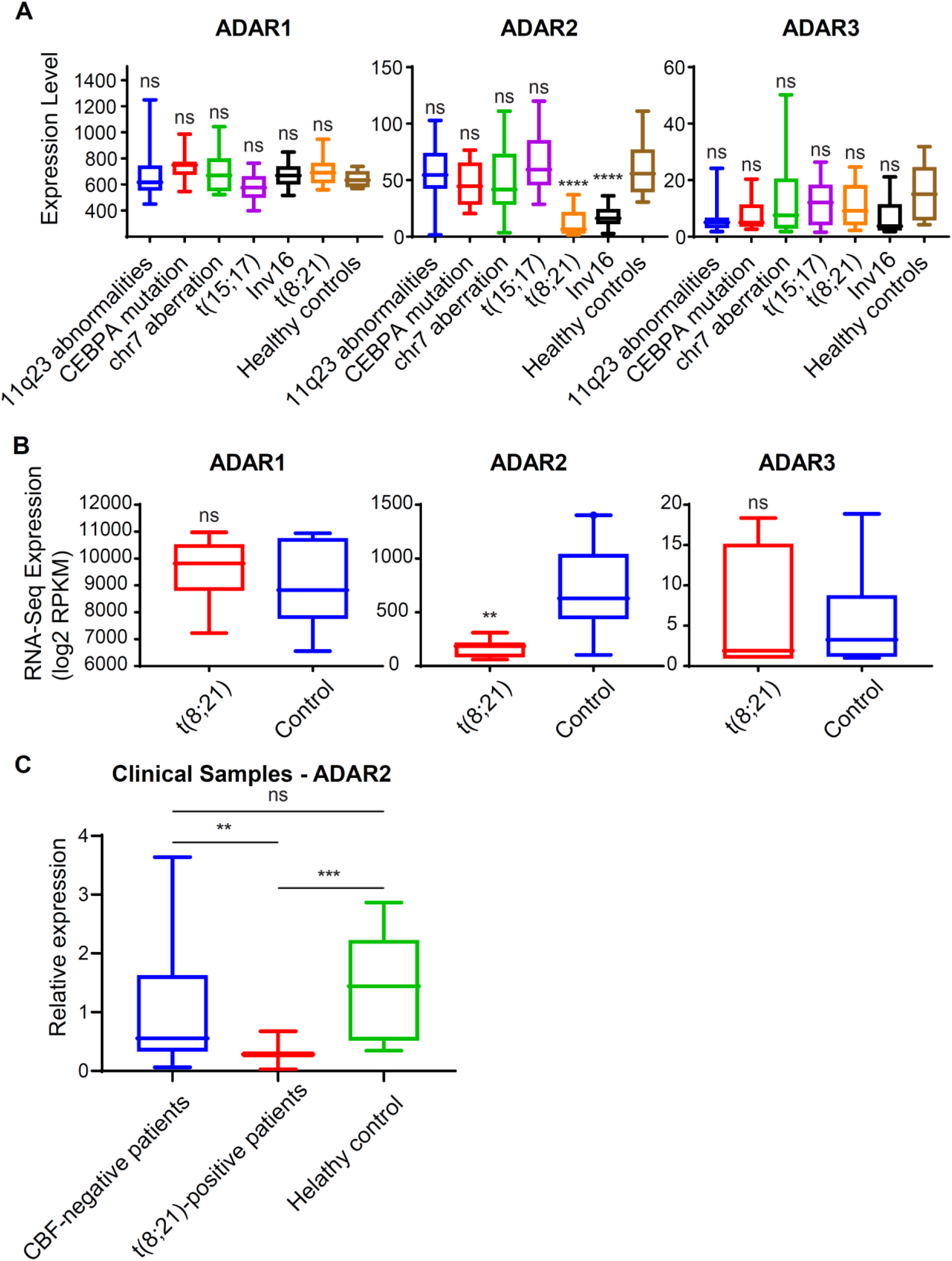
ADAR2 is selectively downregulated in CBF AMLs. **A)** Expression levels of ADAR1, ADAR2, and ADAR3 in healthy donors (n = 8) and different subtypes of AML patients (n=108) (****, *p* < 0.0001, *n*.*s*., not significant; two-tailed Student’s *t*-test). Gene expression analysis was conducted a publicly available Affymetrix microarray dataset downloaded from the GEO database (GSE1159)^19^. The level of expression of a particular gene is reflected by the intensity of hybridization of labelled messenger RNA (mRNA) to gene-specific probe sets (10 to 20 oligonucleotides per gene)^19^. **B)** Expression of *ADAR1, ADAR2, ADAR3* in t(8;21) AML patients (n=7) and the control group including AML patients without t(8;21) and inv16 (n=10), from TCGA. (**, *p* < 0.01, *n*.*s*., not significant; two-tailed Student’s *t*-test) **C)** Quantitative PCR (qPCR) analysis of *ADAR2* transcript level in leukemic blasts isolated from t(8;21)-positive (n=11) and CBF-negative (n=24) AML patients as well as CD34-positive cells isolated from bone marrow samples of healthy individuals (n=16). Data are presented as the mean ± SD of technical triplicates from a representative experiment. (**, *p* < 0.01, ***, *p* < 0.001, *n*.*s*., not significant; two-tailed Student’s *t*-test.)

### RUNX1-ETO and its truncated variant AE9a demonstrate dominant negative effects on *ADAR2* transcription in t(8;21) AML

Next, we investigated the mechanism underlying the downregulation of ADAR2 in t(8;21) AML. It is reported that the RUNX1-ETO fusion protein works as a dominant-negative competitor of RUNX1 against RUNX1-mediated gene expression^32-36^, it is possible that RUNX1 may transcriptionally activate *ADAR2* expression, and such a regulation may be hindered by RUNX1-ETO which outcompetes RUNX1 for binding to the *ADAR2* promoter. To this end, 5000 bp upstream of the transcription start site (TSS) of *ADAR2* and found four putative RUNX binding sites (TGTGGT) (**Figure 2A**). To investigate whether RUNX1 and/or RUNX1-ETO bind to these sites, chromatin immunoprecipitation was conducted in Kasumi-1 cells, a human t(8:21) AML cell line, using anti-RUNX1 (to pull down wildtype RUNX1) or anti-ETO ^37^) antibody, followed by qPCR analysis of three different regions (R1, R2, and R3) **(Figure 2A**). As a result, both RUNX1 and RUNX-ETO could bind to the distal regulatory region of *ADAR2*, as evident from the observation that R1 and R2 regions of *ADAR2* gene showed approximately 100- and 20-fold enrichment in both RUNX1 and RUNX1-ETO pulldown samples compared to the IgG counterparts, respectively (**Figure 2B**). We further determined whether these RUNX1 sites are essential for transcription activation of *ADAR2*. We generated reporter constructs by inserting different DNA fragments (A1-A4) upstream of *ADAR2* TSS. A dramatic drop in the luciferase signal was observed in Kasumi-1 cells when the RUNX site 1 and site 2 were deleted in the A3 fragment (**Figure 2C**), indicating that site 1 and site 2 are indeed essential for the transcriptional activation of *ADAR2* gene.

**Figure 2.**
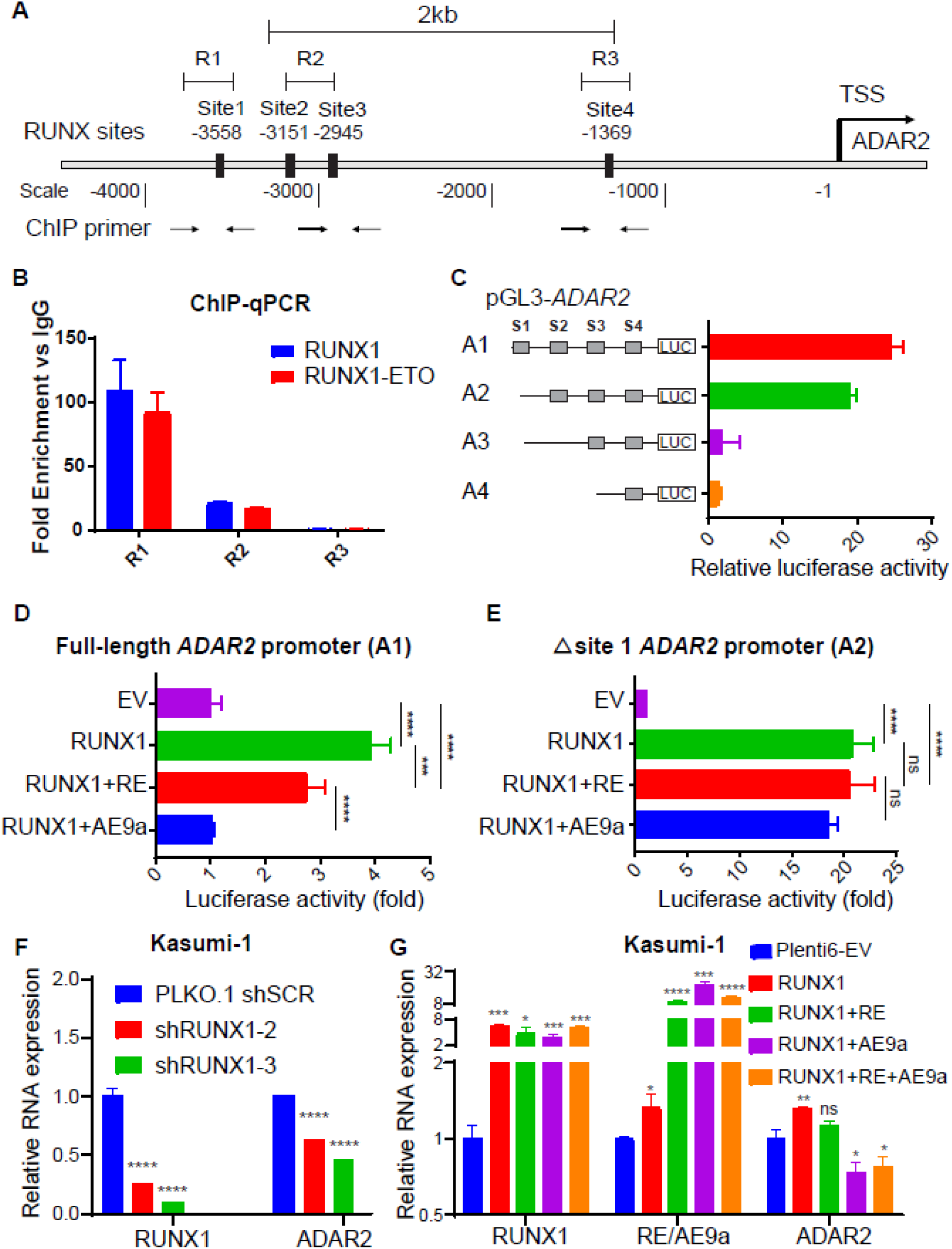
RUNX1-ETO and its truncated variant AE9a demonstrate dominant negative effects on *ADAR2* transcription in t(8,21) AML. **A)** Schematic diagram of the RUNX1 binding sites along the 4kb region upstream of the transcription start site (TSS) of *ADAR2* gene. Number indicates the position with respect to the *ADAR2* TSS which is at position -1. Black bars indicate four putative RUNX1 binding sites. The locations of primers used for ChIP-qPCR experiments are indicated by black arrows. Primers were designed to amplify R1, R2, and R3 regions which cover site 1, site 2 & 3, and site 4, respectively. **B)** ChIP-quantitative PCR (ChIP-qPCR) analysis of the binding of RUNX1 or RUNX1-ETO protein to the indicated regulatory region (R1, R2, and R3) upstream of the TSS of *ADAR2* gene in Kasumi-1 cells, using anti-RUNX1 or anti-ETO antibody respectively. IgG was used as a negative control. Data are presented as the mean ± SD of technical triplicates from a representative experiment. The anti-RUNX1 antibody recognizes the N-terminal portion of RUNX1 protein; while the anti-ETO antibody was used for immunoprecipitation to specifically pull down RUNX1-ETO in Kasumi-1 cells which do not express wildtype ETO protein. **C)** Bar charts demonstrate the luciferase activities associated with each of the indicated sequences upstream of the TSS of *ADAR2*. HEK293T cells were transfected with each of the indicated reporter constructs containing A1, A2, A3, or A4 fragment. S1-S4, site 1, 2, 3 and 4. Luciferase activity of each reporter construct is normalized to the pGL3 empty vector control and defined as ‘Relative luciferase activity’. Data are presented as the mean ± SD of three independent experiments. **D)** Bar charts demonstrate the luciferase activities associated with the A1 fragment in HEK293T cells that were transfected with *RUNX1* alone, or *RUNX1* together with *RUNX1-ETO* or *AE9a* (RUNX1, RUNX1+RE, or RUNX1+AE9a respectively). To calculate the fold change of the luciferase activity, luciferase activity associated with the A1 fragment detected in the indicated group was divided by that of the empty vector (EV) control. Data are presented as the mean ± SD of three independent experiments. **E)** Bar charts demonstrate the luciferase activities associated with the A2 fragment in HEK293T cells that were transfected with *RUNX1* alone, or *RUNX1* together with *RUNX1-ETO* or *AE9a* (RUNX1, RUNX1+RE, or RUNX1+AE9a respectively). Data are calculated and presented using the same method as described in **D**). Data are presented as the mean ± SD of three independent experiments. **F)** Semi-quantitative PCR (qPCR) analysis of *RUNX1, RUNX1-ETO/AE9a*, and *ADAR2* mRNA expression in Kasumi-1 cells, upon shRNA-mediated knockdown of *RUNX1* (shRUNX1-2 and shRUNX1-3). The relative expression of each gene in the indicated group of cells was calculated by the formula 2^™ΔCT^ (ΔCT = CT_(gene)_ –CT_(β-actin)_) and then normalized to the scramble shRNA control counterpart (PLKO.1 shSCR, defined as 1.0). Data are presented as mean ± SD. of three independent experiments (****, *p* < 0.0001, *n*.*s*., not significant; two-tailed Student’s *t*-test.) **G)** qPCR analysis of *RUNX1, RUNX1-ETO/AE9a*, and *ADAR2* mRNA expression in Kasumi-1 cells, upon overexpression of empty vector (Plenti6-EV), *RUNX1* alone, *RUNX1* together with *RUNX1-ETO* or *AE9a*, or *RUNX1* together with *RUNX1-ETO* and *AE9a* (RUNX1, RUNX1+RE, RUNX1+AE9a, or RUNX1+RE+AE9a). Data are calculated and presented using the same method as described in **F**) (****, *p* < 0.0001, ***, *p* < 0.001, **, *p* < 0.01, *, *p* < 0.05, *n*.*s*., not significant; two-tailed Student’s *t*-test). Data are presented as the mean ± SD of technical triplicates from a representative experiment.

Next, upon overexpression of *RUNX1* alone (RUNX1) or together with *RUNX1-ETO* (RUNX1+RE) in Kasumi-1 cells, although a significant increase in the luciferase activity of the A1 fragment was detected in both conditions when compared to that of the empty vector control (EV), co-transfection of *RUNX1* and *RUNX1-ETO* was found to repress the luciferase activity of the A1 but not the A2 fragment induced by *RUNX1* overexpression alone (**Figure 2D**). It has been reported that alternative splicing generates a truncated variant of *RUNX1-ETO*, namely AML1-ETO9a (AE9a), containing an additional exon 9a, and RUNX1-ETO and AE9a are co-expressed in most of AML patients with t(8;21) translocation^38^. Co-expression of RUNX1-ETO and AE9a was shown to induce a more aggressive leukemic phenotype with a rapid onset of AML in a retrovirally transduced mouse model^39^. We also examined the combinatory effect of RUNX1-ETO and AE9a on the luciferase activity of the A1 or A2 fragment and found that AE9a has stronger repressive effect on *ADAR2* transcription than that of RUNX1-ETO (**Figure 2D, 2E**). These findings suggested that RUNX1 activates *ADAR2* transcription and expression through binding to the site 1 and 2 at the distal regulatory region of *ADAR2* gene, whereas RUNX1-ETO and its truncated form AE9a may compete with RUNX1 for binding to the RUNX site 1 to suppress *ADAR2* transcription.

Furthermore, we intended to confirm the dominant negative effects of RUNX1-ETO and AE9a on endogenous ADAR2 expression. We first confirmed that endogenous ADAR2 expression could be significantly repressed by specifically knocking down *RUNX1* in Kasumi-1 cells (**Figure 2F**). Next, the co-overexpression of *RUNX1* and *RUNX1-ETO* and/or *AE9a* (RUNX1+RE, RUNX1+AE9a, and RUNX1+RE+AE9a) was found to repress the upregulation of *ADAR2* induced by RUNX1 alone (RUNX1) (**Figure 2G**). In sum, all these results support our hypothesis that RUNX1-ETO and AE9a demonstrate dominant negative effects on *ADAR2* transcription and expression through outcompeting wildtype RUNX1 for binding to the distal regulatory region of *ADAR2* gene.

### Restoration of ADAR2-reguated RNA editing inhibits leukemogenic ability of t(8;21) AML cells

Due to the downregulation of ADAR2, it is not surprising that ADAR2-regulated RNA editome may be suppressed in t(8;21) AML. We queried whether ADAR2 may regulate leukemogenesis of t(8:21) AML cells through its RNA editing function. To this end, the wildtype (ADAR2 WT), catalytically inactive (DeAD mutant^40^), or RNA binding -depleted (EAA mutant^41^) *ADAR2* expression construct was stably transduced in Kasumi-1 cells using a retroviral system. Upon stable overexpression of wildtype ADAR2 but not the DeAD or EAA mutant of ADAR2 in Kasumi-1 cells, a significant reduction in the colony-forming ability of Kasumi-1 cells was observed when compared to the control cells (**Figure 3A** and **3B**). This result suggested that RNA editing function of ADAR2 is required for its suppressive role in leukemogenesis of t(8;21) AML.

**Figure 3.**
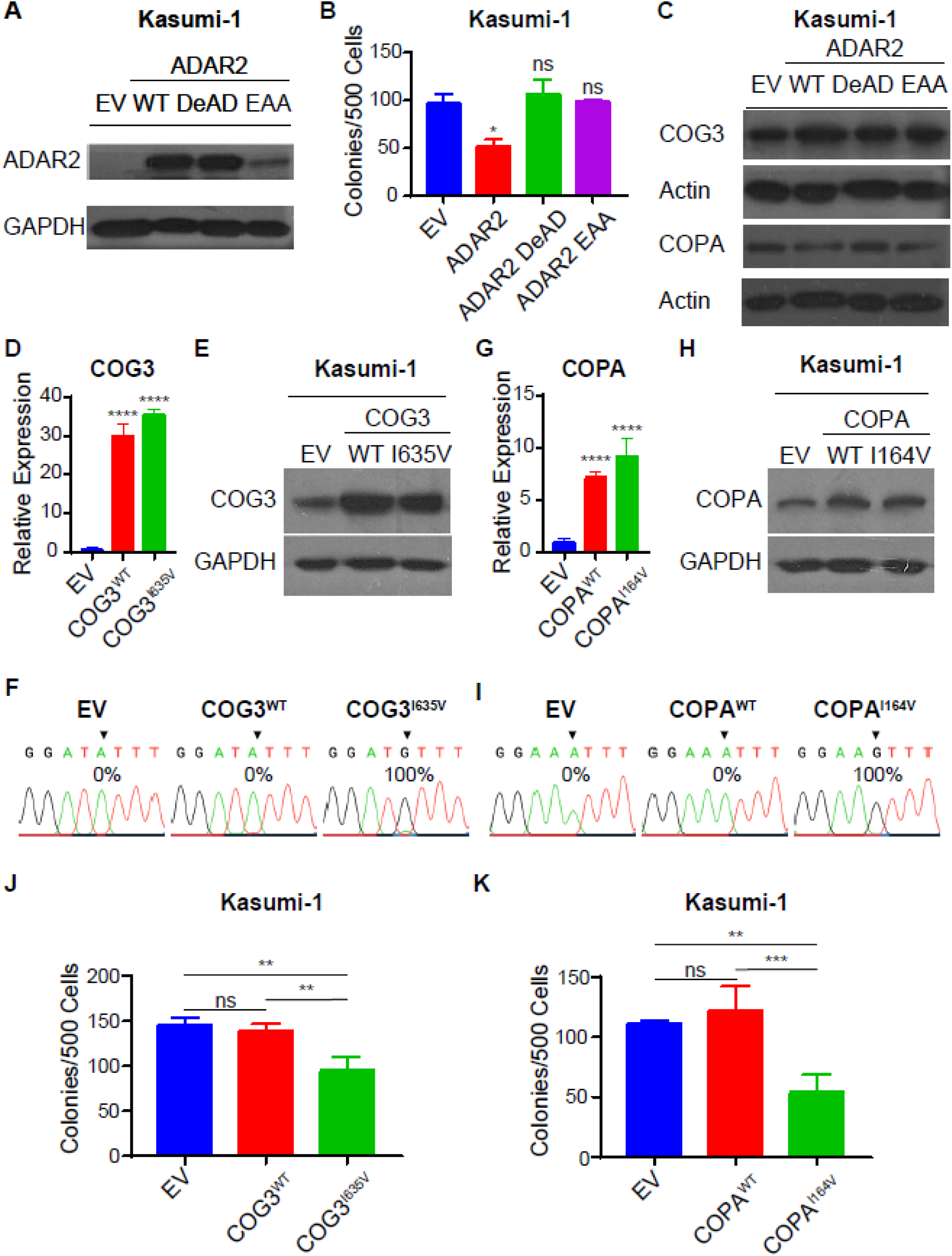
Rescue of ADAR2-repressed clonogenic growth of Kasumi-1 cells through stable overexpression of COPA^I164V^ or COG3^I635V^. **A)** Western blot analysis of ADAR2 protein in Kasumi-1 cells stably overexpressed with the wildtype or mutant form of *ADAR2* by a retrovirus-mediated transduction system. GAPDH was used as a loading control. WT, wildtype ADAR2; DeAD, ADAR2 DeAD mutant; EAA, ADAR2 EAA mutant; EV, empty vector. **B)** Bar chart represents the number of colonies formed by the same group of cells as described in **A**). Data are presented as the mean ± SD of technical replicates from a representative experiment of three independent experiments. (*, *p* < 0.05, *n*.*s*., not significant; two-tailed Student’s *t*-test.) **C)** Western blot analysis of COG3 or COPA protein expression in the same samples as described in **A**). GAPDH was used as a loading control. **D)** qPCR analysis of *COG3* transcript level in Kasumi-1 cells stably overexpressing the wildtype or edited COG3 (COG3^WT^ or COG3^I635V^) or the MSCV-PURO empty vector (EV) control. Data are presented as the mean ± SD of technical triplicates from a representative experiment of three independent experiments. (****, *p* < 0.0001, two-tailed Student’s *t*-test.) **E)** Western blot analysis of COG3 protein level in the same samples as described in **E**). GAPDH was used as a loading control. **F)** Sequence chromatograms illustrate the editing level of *COG3* transcripts in the same samples as described in **D**). The arrow indicates the editing position. **G, H)** qPCR **G**) or western blot **H**) analysis of COPA expression at transcript or protein level respectively, in Kasumi-1 cells stably overexpressing the wildtype or edited COPA (COPA^WT^ or COPA^I164V^) or the MSCV-PURO empty vector (EV) control. Data are presented as the mean ± SD of technical triplicates from two representative experiment of three independent experiments. (****, *p* < 0.0001, two-tailed Student’s *t*-test.) **I)** Sequence chromatograms illustrate the editing level of *COPA* transcripts in the same samples as described in **G-H**). The arrow indicates the editing position. **J, K)** Bar chart represents the number of colonies formed by the same group of cells as described in **D**) and **G, H**). Data are presented as the mean ± SD of technical replicates from one representative experiment from D) and two from G, H) of three independent experiments. (**, *p* < 0.01, *n*.*s*., not significant; two-tailed Student’s *t*-test.)

Next, *COPA* (coatomer subunit α) and *COG3* (Component Of Oligomeric Golgi Complex 3), two reported protein-recoding editing targets regulated by ADAR2^42,43^, were chosen to study whether restoration of expression of the edited protein variant (COPA^I164V^ or COG3^I635V^) could at least partially phenocopy ADAR2-mediated suppression of leukemogenesis of t(8;21) AML. We first confirmed that upon overexpression of the wildtype ADAR2 but not the DeAD or EAA mutant, the editing frequencies of editing sites in *COPA* and *COG3* transcripts were dramatically increased in Kasumi-1 cells (**Supplemental Figure 2A, 2B**), indicating *COPA* and *COG3* are indeed ADAR2 targets in t(8;21) AML cells. Moreover, overexpression of wildtype or mutant form of ADAR2 had no obvious effect on the expression of COPA and COG3 (**Supplemental Figure 2C, 2D** and **Figure 3C**). We next stably expressed the wildtype or edited form of COPA or COG3 (COPA^WT^ and COPA^I164V^, COG3^WT^ and COG3^I635V^) in Kasumi-1 cells (**Figure 3D-3I**). Intriguingly, Kasumi-1 cells constitutionally expressing COPA^I164V^ or COG3^I635V^ but not COPA^WT^ or COG3^WT^ demonstrated significantly lower colony-forming ability compared to the control cells (**Figure 3J, 3K**, and **Supplemental Figure 2E, 2F**). Taken together, these results suggested that restoration of ADAR2-reguated RNA editing inhibits leukemogenic ability of t(8;21) AML cells.

### RNA editing capability of ADAR2 is essential for its repression of leukemogenesis in an AE9a mouse model

It has been reported that co-expression of RUNX1-ETO and its truncated variant AE9a leads to a rapid development of AML in an experimental mouse system^39^, likely due to the impaired transcriptional regulation of RUNX1-ETO target genes^44^. We therefore established an AE9a AML mouse model as reported previously^39^ for further investigation of the role of ADAR2 in t(8,21)-associated leukemogenesis (**Figure 4A**). Of note, *ADAR2* expression was consistently decreased in both bone marrow (BM) cells from recipients following the 1^st^ transplantation, and in that from recipients subjected to serial dilution assay during the 2^nd^ transplantation, suggesting a negative role of ADAR2 in leukemia initiating-potential (**Figure 4B**). To further investigate the function of ADAR2 in leukemogenesis *in vivo*, BM cells from recipients following the 2^nd^ transplantation were transduced with the MSCV-IRES-tdTomato-hADAR2 expression construct or empty vector (**Figure 4C, Supplemental Figure 3A, 3B**). Stably transduced cells were transplanted into sublethally-(6.5Gy) γ-irradiated C57BL/6J mice for peripheral blood (PB) collection and survival monitoring. PB harvested 28 days post-transplantation revealed that white blood cell (WBC) counts backed to the normal region in the *ADAR2* group compared with that in the empty vector group, while red blood cell (RBC) and platelet counts increased slightly in the *ADAR2* group (**Figure 4D**). Moreover, overexpression of *ADAR2* significantly extended mice survival (**Figure 4E**). Wright Giemsa stain confirmed that the majority of BM cells harvested from the *ADAR2* group underwent differentiation, whereas in empty vector group most of them were myeloblast (**Supplemental Figure 3C**). In line with a significant reduction of GFP-positive population in PB from *ADAR2* transduced mice (**Supplemental Figure 3D**), we concluded that ADAR2 plays important function to prohibit leukemogenesis of t(8:21) AML.

**Figure 4.**
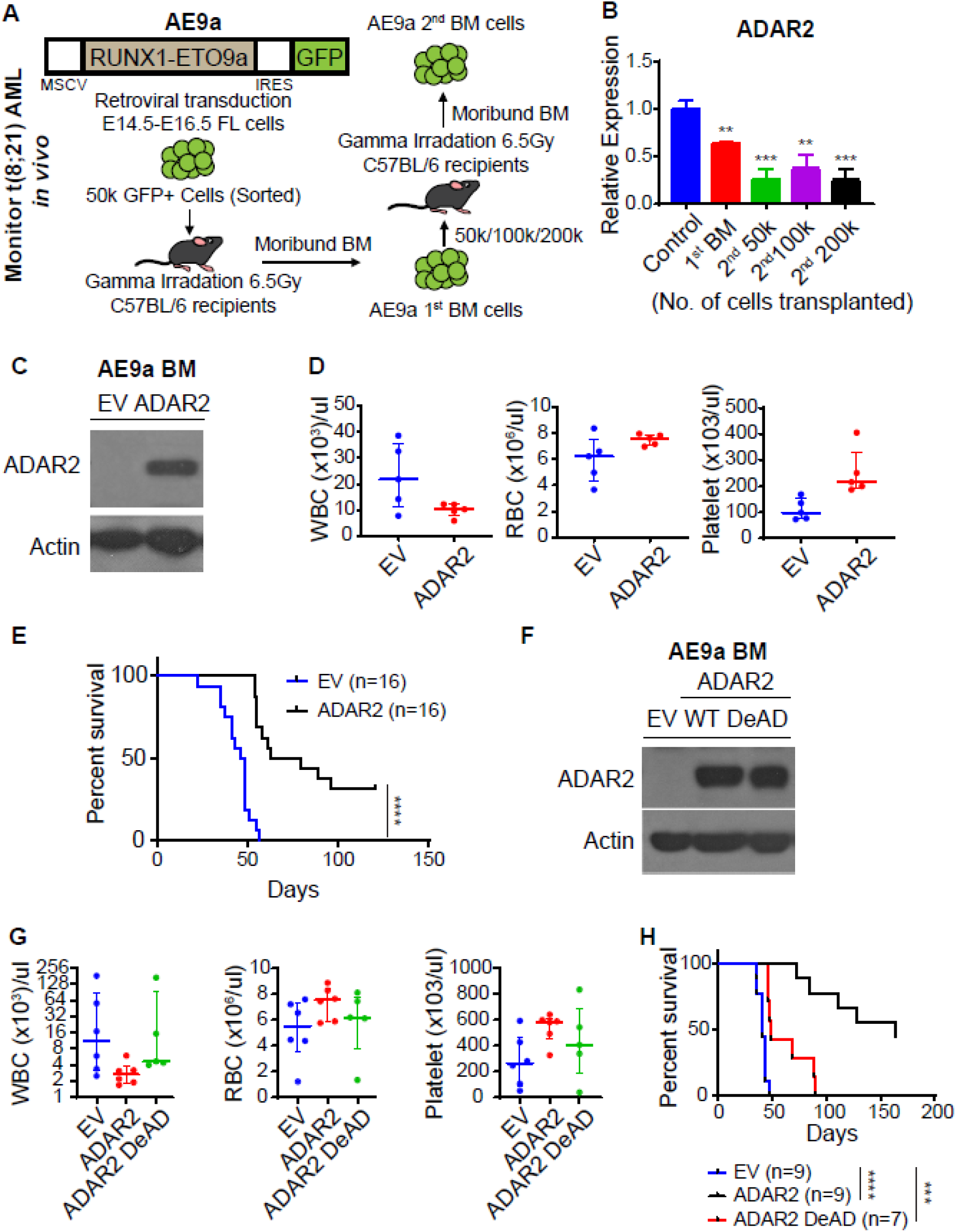
RNA editing capability of ADAR2 is essential for its repression of leukemogenesis in an AE9a mouse model. **A)** Experimental strategy for AE9a bone marrow transplantation mouse model. **B)** qPCR analyses of *ADAR2* transcript in BM cells from moribund mice in the 1^st^ transplantation and in the 2^nd^ transplantation injected with different number of AE9a cells (200k, 100k, 50k). Data are presented as the mean ± SD of technical triplicates from a representative experiment. (**p < 0.01, *p< 0.05, two-tailed Student’s t-test) **C)** Western blot analysis of ADAR2 protein expression in AE9a BM cells stably overexpressing ADAR2 or the MSCV-IRES-tdTOMATO empty vector (EV). **D)** Dot plot represents counts of white blood cell (WBC), red blood cell (RBC), and platelet in the peripheral blood from recipients at 28 days post-transplantation of AE9a AML cells stably overexpressing ADAR2 or the MSCV-IRES-tdTOMATO empty vector (EV). n=5 in each group. **E)** Kaplan–Meyer survival curve of recipients transplanted with 50,000 AE9a AML cells stably overexpressing ADAR2 or MSCV-IRES-tdTOMATO empty vector (EV). n=16 in each group. Statistical analysis is performed using Log-rank (Mantle-Cox) test. (***p < 0.001). **F)** Western blot analysis of ADAR2^WT^ or ADAR2^Mut^ protein expression in BM cells from the same recipients as described in E). Actin was used as a loading control. **G**. Dot plot represents counts of white blood cell (WBC), red blood cell (RBC), and platelet in the peripheral blood from recipients at 45 days post-transplantation of AE9a AML cells stably overexpressing of ADAR2^WT^ (n=6), ADAR2^Mut^ (n=5), or MSCV-IRES-tdTOMATO empty vector (n=6). **H)** Kaplan–Meyer survival curve of recipients transplanted with 50k AE9a AML cells stably overexpressing of ADAR2^WT^ (n=9), ADAR2^Mut^ (n=7), or MSCV-IRES-tdTOMATO empty vector (EV) (n=9). Statistical analysis is performed using Log-rank (Mantle-Cox) test. (****p < 0.0001, ***p < 0.001.)

To further explore whether RNA editing function of ADAR2 is required for ADAR2-mediated leukemogenesis inhibition, we overexpressed wildtype ADAR2 (ADAR2^WT^) or the DeAD mutant (ADAR2^Mut^) in BM cells from the recipients following 2^nd^ transplantation using a retrovirus-mediated transduction system. After verifying of the transduction efficiency (**Figure 4F, Supplemental Figure 3E**), cells were transplanted back into sub-lethally (6.5Gy) γ-irradiated C57BL/6J mice for PB collection and survival monitoring. Comparing with empty vector group, lower WBC count, and higher RBC or platelet count was detected in wildtype *ADAR2* group, but not in *ADAR2 DeAD* or *ADAR2 EAA* groups. (**Figure 4G**). Furthermore, although approximately 50% of ADAR2^WT^ mice survived over 150 days, all ADAR2^Mut^ recipients died within 100 days, while all control mice (EV) died within 50 days (**Figure 4H**). Consistently, Wright Giemsa stain revealed that loss of RNA editing ability impacted ADAR2-mediated BM cell differentiation (**Supplemental Figure 3F**). Higher percentage of AE9a (GFP)-positive cells were also detected in BM cells of ADAR2^Mut^ recipients than the ADAR2^WT^ counterparts (**Supplemental Figure 3G**). Taken together, these data indicate that ADAR2 supresses leukemogenesis of t(8:21) AML via RNA editing *in vivo*.

## Discussion

A-to-I RNA editing is one of the most common posttranscriptional RNA modification processes that change the DNA coding in the mammalian transcriptome. Enzymes catalysing this process include ADAR1, ADAR2, ADAR3^3^. ADARs demonstrate specific expression patterns in different tissues and environments^45,46^. While the dysregulation of A-to-I RNA editing is implicated in human diseases including multiple cancers^6,8,10-12^, roles of RNA editing in leukemia, particularly ADAR2 and ADAR2-mediated RNA editing, remain largely unexplored.

In this study, utilizing both clinical samples and mouse models, we reveal a previously undescribed role of ADAR2-mediated RNA editing in leukemogenesis. Initiating with a serial of analyses with different cohorts of patients with AML, we found that *ADAR2* is specifically down-regulated in CBF AML patients, whereas no obvious changes in the expression levels of *ADAR1* and *ADAR3* among different subtypes of AML. Mechanistically, we found that the wildtype RUNX1 activates expression of *ADAR2* transcriptionally through two distal RUNX binding sites (1 and 2) upstream of the TSS of *ADAR2* gene, whereas RUNX1-ETO and its truncated variant AE9a suppress RUNX1-mediated activation of *ADAR2* transcription in a dominant-negative manner, likely by outcompeting RUNX1 for binding to the site 1. Due to the fact that ADAR2 can suppress the development and/or progression of solid tumours such as HCC and CRC through RNA editing^42,47^, it prompted us to study whether ADAR2 may lead to leukemogenesis of t(8;21) AML via its RNA editing function. Indeed, as evident from our cell line-based studies and mouse experiments, ADAR2 prohibits leukemogenesis of t(8;21) AML dependent of its RNA editing ability. It is an exciting discovery on the role of RNA editing in t(8;21) AML. Previous studies on this AML subtype mainly focus on the changes of gene expression patterns or chromatin status^48^; and the gap between AML and RNA editing is still open. Our results showed that restoration of ADAR2 level in AE9a-positive BM cells significantly prolongs survival of recipients, whereas RNA editing-deficient form has very limited rescue efficiency. This finding clearly indicates that ADAR2 and it mediated RNA editing is of biological importance to inhibit leukemogenesis in t(8;21) AML. Of note, the limited rescue effect of the editing-deficient ADAR2^Mut^ is likely to be attributed to the RNA editing-independent functions of ADAR2. For example, ADARs can regulate microRNA maturation through interaction with Dicer ^49^. ADAR2 enhances target mRNA stability by limiting the interaction of RNA-destabilizing proteins with their cognate substrates^50^. In general, our study sheds a new light on a novel role of ADAR2 in suppressing leukemogenesis of t(8;21) AML via RNA editing.

In line with other studies, our study further emphasizes the importance of RNA metabolism in AML and opens a new avenue for the development of RNA therapies for AML. Functions of RNA processing steps such as splicing, polyadenylation, RNA modifications and RNA editing in AML remain completely unknown until recent years. For example, leukemogenesis through reducing of N6-methyladenosine (m^6^A) RNA modification by FTO (Fat Mass and Obesity-associated protein) is firstly discovered in 2017^51^. Since then, numbers of studies from several groups described the importance of m^6^A modification in normal hematopoiesis and AML, suggesting m^6^A RNA modification as a potential process for AML therapy^52-59^. Similar to m^6^A RNA modification, A-to-I RNA editing is the most prevalent type of RNA editing in mammals. Our result revealed that restoration of ADAR2-mediated RNA editing of COG3^I635V^ and COPA^I164V^ could inhibit colony-forming of Kasumi-1 cells. Together with previous findings that COG3^I635V^ could increase cell viability in multiple normal human cell lines and drug sensitivity including MEK inhibitors^9^ and COPA^I164V^ functions as a dominant negative form which represses the oncogenic function of the unedited and wildtype COPA (COPA^WT^)^42^, our results suggest a novel therapeutic strategy by restoring edited RNAs. Notably, in the present study, only two ADAR2-regulated editing targets were tested in cell culture-based experiments. To perform a transcriptome-wide identification of ADAR2 target genes in different cell populations/types, single cell RNA-seq will be required, and the functional importance of potential editing targets needs to be verified by conducting rescue experiments in mouse AML model or pre-clinical AML model such as AML patient-derived organoids (PDOs).

Another interesting finding is the involvement of CBF complex in regulation of *ADAR2* transcription and expression. Expression of *ADARs* is commonly dysregulated in multiple cancer types. For instance, expression of *ADAR1* is dysregulated in esophageal squamous cell carcinoma (ESCC), HCC, gastric cancer, cervical cancer, lung cancer, and breast cancer, and disruption of *ADAR2* expression was detected in gastric cancer, HCC, ESCC, and glioblastoma^6,7,12,13,60-67^. However, the regulatory mechanisms leading to dysregulation of *ADARs* expression remain elusive. In our study, we found that RUNX1-ETO and AE9a demonstrate dominant negative effects on *ADAR2* transcription and expression through outcompeting wildtype RUNX1 for binding to the distal regulatory region of *ADAR2* gene. Since both RUNX1 and ADAR2 express in multiple organs, it is possible that such regulatory mechanism commonly presents in other tissues. On the other hand, besides t(8;21) AML, *ADAR2* was also significantly downregulated in the other type of CBF leukemia-inv(16) which generates *CBFβ-MYH11* fusion gene. As the CBFβ-MYH11 fusion protein retains the ability to bind RUNX1 with increased affinity as compared to CBFβ, thereby sequestering RUNX proteins from their target genes^68-71^. This may account for the downregulation of *ADAR2* in inv(16) AML patients and also provides a hint on the regulation of *ADAR2* expression by CBF complex, which remains for our further investigation. Of note, changes in chromatin accessibility regulates transcription of *ALKBH5*, an important m^6^A demethylase required for maintaining leukemia stem cells (LSCs) function^72^. Due to the fact that chromatin status, such as histone modifications and chromatin interactions, plays important role in hematopoiesis^73,74^, another aspect to consider is that the regulation of *ADAR2* expression through dynamic changes of chromatin status. In sum, our findings shed new light on future studies of dysregulated *ADAR2* transcription and expression as well as their RNA editing-dependent or independent biological implications in multiple diseases including cancers.

## Acknowledgement

We thank Michelle, Mok Meng Huang for assistance with flow cytometry experiments and Wright Giemsa stain and thank Dr. Sudhakar Jha for providing the pMSCV-flag-puro plasmid. This research was supported by the Singapore Ministry of Health’s National Medical Research Council (NMRC), the Singapore National Research Foundation (NRF), the Singapore Ministry of Education (MOE) under its Research Centres of Excellence initiative (RCE), and the U.S. National Institutes of Health (NIH) with following detail: Singapore Translational Research (STaR) Investigator Award, NIH Grants 1R35CA197697 and P01 HL131477 to D.G.T., MOE Tier 2 Grants MOE2018-T2-1-005 and MOE2019-T2-2-008, NMRC Clinician Scientist-Individual Research Grant (CS-IRG: MOH-CIRG18nov-0007, Project ID: MOH-000214) to L.C., NMRC Open Fund-Young Individual Research Grant MOH-OFYIRG20nov-0011 and MOE Research Scholarship Block (RSB) Grant N-171-000-019-001 to Q.Z., as well as MOE Tier 3 Grant (MOE2014-T3-1-006) to D.G.T. and L.C.

## Authorship Contributions

D.G.T. and L.C. conceived and co-supervised the study with Q.Z. D.G.T., L.C., M.G., T.H.M.C., and Q.Z. designed the experiments. M.G. and T.H.M.C. performed all experiments with input from Y.S. and V.H.E.N. H.Y. and O.A. conducted all bioinformatics analyses. Z.H.T. and V.H.E.N. assisted with mouse experiments. W.J.C. and M.O. provided the leukemia clinical samples. D.G.T., L.C, Q.Z and M.G. wrote the manuscript with input from T.H.M.C and X.C.

