## Supplemental information for "ADAR2-repressed RNA editing: a novel mechanism contributing to t (8:21) AML leukemogenesis"

**Affiliations:**

Supplemental information

1. Supplemental materials and methods

2. Supplemental figures and figure legends

3. References

#### Supplemental materials and methods

##### Cell lines

HEK293T (ATCC, cat.no. CCL-11268), Bosc 23 (ATCC, cat.no. CRL-11270), GP2-293 (Clontech, cat.no. 631530) cells were cultured in DMEM high glucose medium (Biowest, cat.no. L0104) supplemented with 10% FBS (Hyclone, cat.no. SH30071). Kasumi-1 (ATCC, cat.no. CRL-2724) cells was maintained in RPMI high glucose medium (Biowest, cat.no. L0500) supplemented with 20% FBS. Cells were incubated at 37 °C in a humidified incubator containing 5% $CO_2$. All cells used in this study were regularly authenticated by morphologic observation.

##### RNA isolation, RT-PCR and RT-qPCR

RNA isolation was performed using Trizol reagent (Invitrogen, cat.no. 15596026) and RNeasy Mini Kit (Qiagen , cat.no. 74104) according to the manufacturer's instructions. Reverse transcription reaction was performed using SuperScript VILO cDNA Synthesis Kit (Invitrogen cat.no. 11754-250) according to the manufacturer's instructions. qPCR was performed using GoTaq® qPCR Master Mix (Promega cat.no. A6002) on the QuantStudio 3 or 5 Real-Time PCR Systems (ThermoFisher Scientific) according to the manufacturer's instructions. Sequence of qPCR primers were listed in Supplemental Table 1.

##### Plasmid construction

To generate lentiviral expression constructs of *RUNX1*, *RUNX1-ETO*, or *AE9a*, full-length *RUNX1*, *RUNX1-ETO*, or *AE9a* were amplified from their overexpressing construction using

Q5® High-Fidelity DNA Polymerase (NEB, cat.no. M0491L) and cloned into pLenti6/V5-TOPO vector through XhoI and SacII restriction sites.

To clone *ADAR2* or *ADAR2* mutants into pMSCV-IRES-tdTomato vector, cDNA of *ADAR2*, *ADAR2 DeAD*, or *ADAR2 EAA* were amplified from their overexpression plasmid[1-3] using Q5® High-Fidelity DNA Polymerase and cloned into empty vector using by "In-Fusion Cloning" strategy.

The full-length of wild type *COPA* cDNA were amplified from its overexpression plasmid[4] using Q5® High-Fidelity DNA Polymerase and cloned into pMSCV-flag-puro vector from Dr. Jha through MluI and NcoI restriction sites. The full-length of wild type *COG3* cDNA were amplified from its into pLenti6/V5-TOPO overexpression plasmid we generated before using Q5® High-Fidelity DNA Polymerase and cloned into pMSCV-flag-puro vector through MluI and NcoI restriction sites. *COG3 I635V* and *COPA I164V* edited forms were generated through PCR-directed mutagenesis using their full-length constructs.

To specifically knockdown *RUNX1*, shRNAs against *RUNX1* were designed following RNA consortium guidelines (http://www.broadinstitute. org/rnai/public/). To specifically target *RUNX1* but not *RUNX1-ETO* or *AE9a*, *RUNX1* shRNAs were designed to against c-terminal of *RUNX1* without overlapping on *RUNX1-ETO* or *AE9a*. shRNAs were cloned into the pLKO.1-puro vector following Addgene pLKO.1 protocol (http://www.addgene.org/tools/protocols/plko/).

For luciferase assay, different lengths of human *ADAR2* promoter region containing different RUNX site portions was PCR amplified from a BAC clone (RP11-280L8) and cloned into pGL3-LUC firefly luciferase reporter vector using MluI and BglII restriction cutting sites

(A1: -3,625 bp to -1 bp from *ADAR2* TSS; A2: -3,179 bp to -1 bp from *ADAR2* TSS; A3: -2,965 bp to -1 bp from *ADAR2* TSS; A4: -1,500 bp to -1 bp from *ADAR2* TSS, Figure 2C).

**Lentiviral Expression System**

To overexpressing different proteins or to knocking down either RUNX1 or RUNX1-ETO in Kasumi-1 cells, lentiviral expression system is utilized. In brief, 2 µg VSV-G, 4 µg delta 8.9, and 4 µg expression constructs were co-transfected into 293T cells cultured in T75 flask for virus generation. Supernatant containing lentivirus was harvested and filtered using 0.45 µm syringe filter (Sartorius, cat.no. 16537) at 48 and 72 hours after transfection. These two portions of supernatant were mixed and aliquoted for virus transduction or stored in -80°C for subsequent usage. For transduction, 0.6 M Kasumi-1 cells were resuspended in lentivirus containing medium plus 10 µg/ml polybrene followed by centrifuging at 2,400 RPM, 32°C for 3 hours. After discarding supernatant, fresh medium was added into the transduced cells. At 72 hours after transduction, cells were collected for subsequent analysis.

**Establishment of Kasumi-1 stable cell lines**

To generate Kasumi-1 cells stably expressing *ADAR2*, *ADAR2* mutants, wild-type or edited forms of *COPA* and *COG3*, 4.5 µg VSV-G and 4.5 µg expression constructs were co-transfected into GP2-293 cells cultured in T75 flask for virus generation. Supernatant containing lentivirus was harvested and filtered using 0.45um syringe filter (Sartorius, cat.no. 16537) at 48 and 72 hours after transfection. These two portions of supernatant were mixed and aliquoted for virus transduction or stored in -80°C for subsequent usage. Virus transduction was performed using RetroNectin® Recombinant Human Fibronectin Fragment (Takara, cat.no. T100A/B) according to the manufacturer’s instructions (with centrifugation). At 72 hours after virus transduction, cells were selected using puromycin (1 µg/ml) for 72 hrs or by FACS sorting.

**Luciferase reporter assays**

The firefly luciferase reporter gene in the pGL3 vector is driven by human *ADAR2* promoter regions containing different RUNX site portions (A1, A2, A3, and A4). As an internal control plasmid for co-transfections, the pRL-null construct encoding a Renilla luciferase gene (Promega, cat.no. E2231) was used. Firefly and Renilla luciferase activities were determined 24 hours post transfection with the dual-luciferase reporter assay system (Promega, cat.no. E1910). To determine the transcription regulation effect of different RUNX1 sites (Figure 2C), after normalization against internal control Renilla luciferase reading in every group, firefly luciferase readings from cells transfected with A1, A2, A3, or A4 were calculated as fold differences against the activity obtained from cells transfected with empty pGL3 vector. To investigate the regulatory mechanism of RUNX1 and RUNX1-ETO on *ADAR2* expression (Figure 2D, 2E), pGL3 vector containing full-length ADAR2 promoter (A1) or truncated ADAR2 promoter (A2) and pRL-null plasmid were co-transfected with empty vector or expression constructs of RUNX1, RUNX1-ETO, or AE9a as indicated in the figure. Firefly luciferase readings were normalized against internal control Renilla luciferase and calculated as fold differences against the activity obtained from cells transfected with empty vector.

**Western-blot analysis**

After PBS washing, cell pellet was resuspended in PBS to a concentration of 4 M per ml. Equal amount of 2x SDS loading dye (4% SDS, 30% glycerol, 2% 2-mercaptoethanol, 0.08% bromphenol blue, 0.1 M Tris HCl, pH 6.8) was added into cell suspension followed by incubation at 97 $^{0}$C for 10 mins to lysate cells. Proteins were separated on 8% SDS-PAGE gels. Immunoblots were incubated with primary antibody overnight at 4 °C, followed by a secondary horseradish peroxidase (HRP)-conjugated antibody (1:5000 dilution) at room temperature for 1 hour. Details on the antibodies used and dilutions are described in Supplemental Table 2.

**Chromatin immunoprecipitation followed by qPCR (ChIP-qPCR)**

Cells were fixed with 1% formaldehyde for 15 mins at RT with rotation. Then formaldehyde was neutralized by adding 2.5 M glycine to a final concentration of 0.125 M and rotating for 5 mins at RT. After washing with PBS, cells were lysed with ChIP SDS lysis buffer (100 mM NaCl,50 mM Tris-Cl pH8.0, 5 mM EDTA, 0.5% SDS, and 1x protease inhibitor (Roche, cat.no. 11836145001) supplied freshly), and then stored at -80 $^{0}$C until further processing. Nuclei were collected by spinning down at 13,000 rpm for 10 mins. The nuclear pellet was re-suspended in IP solution (2 volume ChIP SDS lysis buffer plus 1 volume ChIP triton dilution buffer (100 mM Tris-Cl pH8.6, 100 mM NaCl, 5 mM EDTA, 5% Triton X-100), and 1x protease inhibitor (Roche, cat.no. 11836145001) supplied freshly) for sonication using a Bioruptor® Pico (Diagenode) to obtain 200 bp to 500 bp DNA fragments. After spinning down to remove debris, sonicated chromatin was pre-cleared by adding 30 µl washed dynabeads protein A/G (Invitrogen, cat.no. 10002D/10004D) and rotated at 4 $^{0}$C for 4 hrs. Pre-cleared chromatin was incubated with antibody pre-bound dynabeads protein A/G overnight at 4 $^{0}$C. Next day, magnetic beads were washed through the following steps: buffer 1 (150 mM NaCl, 50 mM Tris-Cl, 1 mM EDTA, 5% sucrose, 0.02% $NaN_3$, 1% Triton X-100, 0.2% SDS, pH 8.0) two times; buffer 2 (0.1% deoxycholic acid, 1 mM EDTA, 50 mM HEPES, 500 mM NaCl, 1% Triton X-100, 0.02% NaCl, pH 8.0) two times; buffer 3 (0.5% deoxycholic acid, 1 mM EDTA, 250 mM LiCl, 0.5% NP40) two times; TE buffer one time. After removing TE buffer, beads were incubated with 120 µl TE buffer containing 1% SDS and 100 µg Recombinant Proteinase K (Invitrogen, cat.no. AM2548) at 65 $^{0}$C overnight for reverse crosslinking. Next day, DNA was purified with MinElute PCR

Purification Kit (Qiagen, cat.no. 28006). Enrichment of wild type RUNX1 or RUNX1-ETO on different regions was detected with qPCR. Sequence of qPCR primers are listed in Supplemental Table 1. Details on antibodies used are described in Supplemental table 2.

**Colony formation assay**

The colony formation assay of the stable cell lines was performed using Human Methylcellulose Complete Media (R&D Systems, cat.no. HSC003). Briefly, after PBS washing, 0.3 ml Cell Resuspension Solution, 1,500 cells were added into 3 ml ice-cold methylcellulose complete media and mixed well using 3 ml syringe with 16 G needle. Using the same syringe and needle, only 1.1 ml of the mixture was equally distributed into 35 mm culture plates to avoid generating bubbles. Colony numbers were counted after 7 days.

**RUNX1-ETO9a primary leukemia model**

To generate RUNX1-ETO9a (AE9a) AML mouse model, 1.5ug envelop, 1.5 ug gag-pol, and 6 ug AE9a expression constructs were co-transfected into the BOSC-23 cells cultured in T75 flask for virus generation. Supernatant containing lentivirus was harvested and filtered using 0.45um syringe filter (Sartorius, cat.no. 16537) at 48 and 72 hours after transfection. These two portions of supernatant were mixed and aliquoted for virus transduction or stored in -80°C for subsequent usage. After sorting from fetal liver cells harvested from C57BL/6J embryos at E14.5– E16.5 stage, LSK population was used for virus transduction. Virus transduction was performed using RetroNectin® Recombinant Human Fibronectin Fragment (Takara, cat.no. T100A/B) according to the manufacturer’s instructions (with centrifugation). At 7 days after transduction, fifty thousands of FACS-sorted GPF positive cells were transplanted into sub-lethally (6.5 Gy) γ-irradiated C57BL/6J mice through retro-orbital injection. Moribund mice were euthanized with $CO_2$ and dissected for spleen, vertebrae, femur, tibia, and hip collection. A secondary transplantation was performed through transplanting two hundred thousand bone marrow cells from the first round of transplantation into sub-lethally (6.5 Gy) γ-irradiated C57BL/6J mice through retro-orbital injection. Moribund mice were euthanized with $CO_2$ and dissected for spleen, vertebrae, femur, tibia, and hip collection. The bone marrow cells were stored in -80°C for subsequent usage.

**Serial dilution assay**

Unfractionated BM cells (200, 000, 100,000, 50,000) from leukemic mice in the $1^{st}$ transplantation were transplanted into sub-lethally (6.5 Gy) γ-irradiated C57BL/6J mice through retro-orbital injection. Moribund mice were euthanized with $CO_2$ and dissected for spleen, vertebrae, femur, tibia, and hip collection. Expression level of *ADAR2* in BM cells was examined.

**Rescue of *ADAR2*/*ADAR2* mutant in AE9a mouse model**

Retrovirus overexpressing *ADAR2* or *ADAR2* mutants were generated in the same way as AE9a retrovirus. BM cells from AE9a leukemic mice in the $2^{nd}$ transplantation were used for virus transduction. Briefly, after $CO_2$ euthanasia, bones were collected and crushed using a pestle with ice-cold PBS, and then bone marrow cell suspension were filtered through a 70 µm BD cell strainer. Red blood cells were removed by RBC lysis buffer treatment (5 ml per mouse) for 5 mins followed by wash with 25 ml PBS (with 2% FBS, 2 mM EDTA). Virus transduction was performed using RetroNectin® Recombinant Human Fibronectin Fragment (Takara, cat.no. T100A/B) according to the manufacturer’s instructions (with centrifugation). Transduced cells were cultured in Stemline® II Hematopoietic Stem Cell Expansion Medium (Sigma-Aldrich, cat.no. S0192) supplemented with 5% FBS (Hyclone, cat.no. SH30071), 100 ng/µL murine recombinant SCF (Peprotech, cat.no. 250-03), 6 ng/µL murine recombinant IL-3 (Peprotech, cat.no. 213-13), 10ng/µL murine recombinant IL-6 (Peprotech, cat.no. 216-16), 20 ng/µL murine recombinant IL-11 (Peprotech, cat.no. 220-11) for 7 days. At day 2 after transduction, fifty thousands of FACS-sorted GPF positive cells were transplanted into sub-lethally (6.5 Gy) g-irradiated C57BL/6J mice through retro-orbital injection for Peripheral blood (PB) collection and survival monitoring. PB samples (~100 µl per mouse) were collected in EDTA-coated capillary tube (Drummond Scientific, cat.no. 1-000-800/12) by submandibular venipuncture with 5-mm Goldenrod animal lancets (Braintree Scientific, cat.no. GR5MM). Counts of nucleated cells was performed using a NIHOKODEN auto blood cell counter under Pre-dilute 20 µl mode. Moribund mice were euthanized with $CO_2$ and dissected for spleen, vertebrae, femur, tibia, and hip collection. BM cells harvested from moribund mice were cytospun and stained with Giemsa's azur-eosin-methylene blue solution (Merck, cat.no. 109204).

**Flow cytometry and fluorescence activated cell sorting (FACS)**

To isolate LSK population, fetal liver cells from C57BL/6J embryos at E14.5– E16.5 stage were treated with 5 ml RBC lysis buffer for 5 mins at room temperature. After washing with 25 ml ice-cold PBS (with 2% FBS, 2 mM EDTA), cells were stained described in the supplemental information for 30 mins. Then cells were washed once with PBS and then were resuspended to a final cell concentration of 40 M per ml for sorting on an Arial I FACS sorter (Beckman Coulter). Post sort was performed for double checking of cell populations and for cell number counting. The gating strategy for LSK population is provided in supplemental figure 4.

**ADARs expression profiling in GSE1159 samples**

GSE1159 cDNA microarray data were downloaded from the NCBI Gene Expression Omnibus (GEO) database[5]. The distribution of normalized and log2-scaled expression levels for ADARs was compared by using unpaired Student T test between tumours and healthy control samples. A P value less than 0.05 was regarded as significant up/down-regulation of ADARs between tumours and healthy control samples.

**ADARs expression profiling in TCGA samples**

RNA-seq data (fastq files) of 27 AML patient samples with t(8;21) (n=7), inv16 (n=10), and normal (n=10) karyotype, from TCGA were downloaded from the dbGaP repository, under accession phs000178.v11.p8 (https://www.ncbi.nlm.nih.gov/projects/ gap/cgi-bin/study.cgi?study_id=phs000178.v11.p8). Each sample was processed as follows: Raw reads were aligned to the reference human genome (hg19) by using STAR (v2.5.2a)[6]. The gene expression quantification was performed by using feature Counts (v1.5.0-p3)[7] to obtain raw counts, which were then normalized by dividing with the total number of uniquely mapped reads in the corresponding sample and multiplying by a factor of 100 million. The distribution of normalized and log2-scaled expression levels for ADAR1, ADAR2, and ADAR2 were compared by using unpaired Student T test between t(8;21)/inv(16) and control samples. A P value less than 0.05 was regarded as significant up/down-regulation of ADARs between t(8;21)/inv(16) and control samples.

#### Supplemental Figure 1

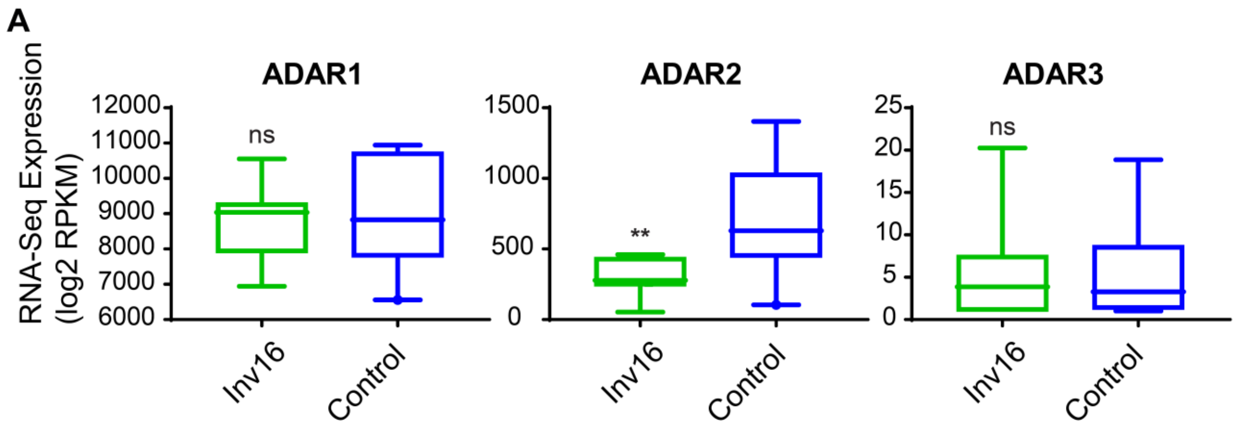


**Supplemental Figure 1. ADAR2 is selectively downregulated in inv16 AML.** Expression of *ADAR1*, *ADAR2*, *ADAR3* in inv16 AML patients (n=10) and the control group including AML patients without t(8;21) and inv16 (n=10), from TCGA. (**, $p < 0.01$, *n.s.*, not significant; two-tailed Student's *t*-test)

### Supplemental Figure 2

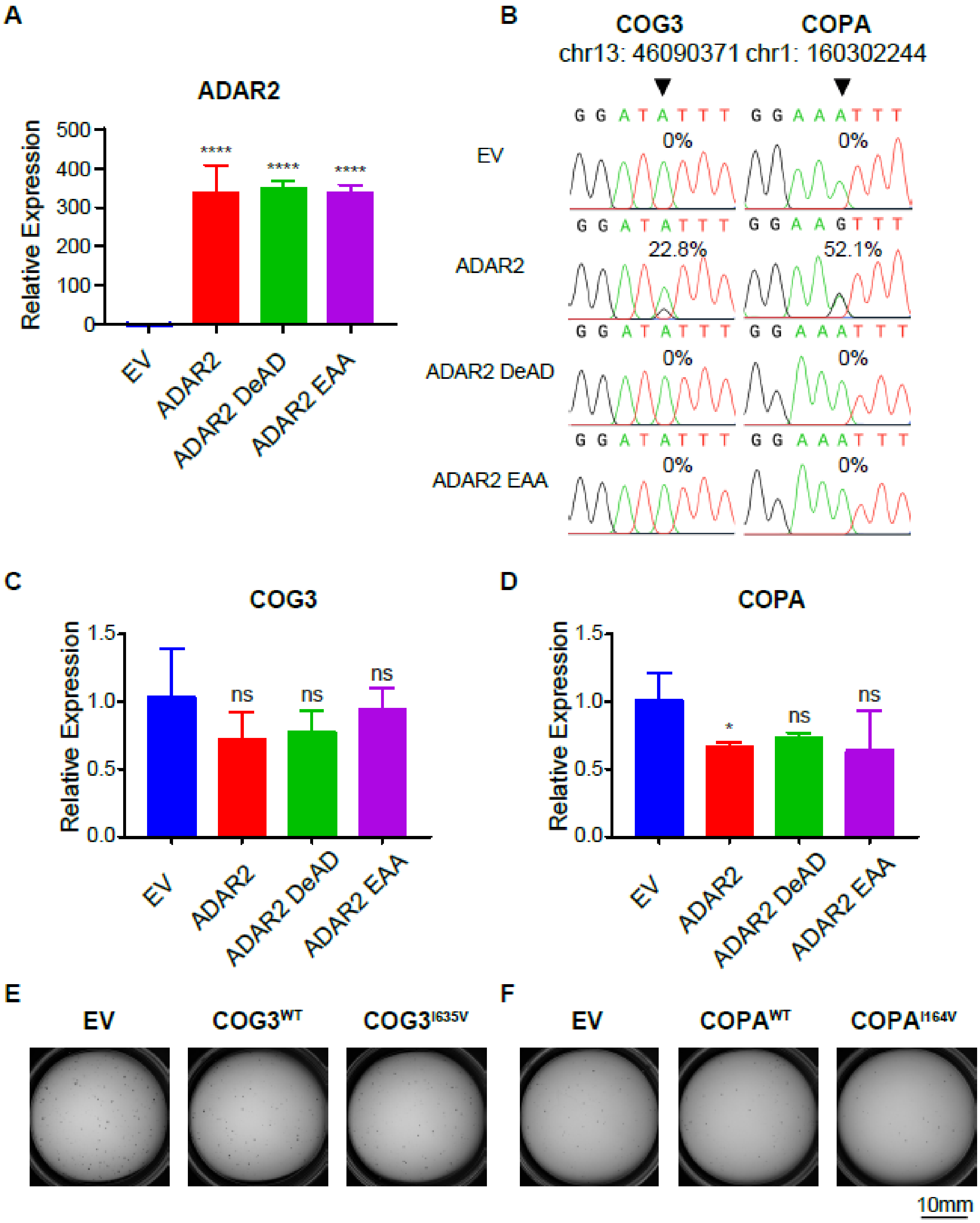

**Supplemental Figure 2. Rescue of ADAR2-repressed clonogenic growth of Kasumi-1 cells through stable overexpression of COPA$^{I164V}$ or COG3$^{I635V}$.**

**A)** qPCR analysis of *ADAR2* transcripts in the same groups of cells as described in Figure 3A. Data are presented as the mean ± SD of technical triplicates from a representative experiment of three independent experiments. (****, $p < 0.0001$, two-tailed Student's *t*-test.)

**B)** Sequence chromatograms showing the editing levels of *COG3* and *COPA* transcripts in the same groups of cells as described in Figure 3A. The arrow indicates the editing position.

**C, D)** qPCR analyses of *COG3* **C**) or *COPA* **D**) transcript in the same groups of cells as described in Figure 3A. Data are presented as the mean ± SD of technical triplicates from a representative experiment. (***, $p < 0.001$, **, $p < 0.01$, *n.s.*, not significant; two-tailed Student's *t*-test.)

**E, F)** Representative images for the colony-forming assay conducted in the same groups of cells as described in Figure 3E-3I.

#### Supplemental Figure 3

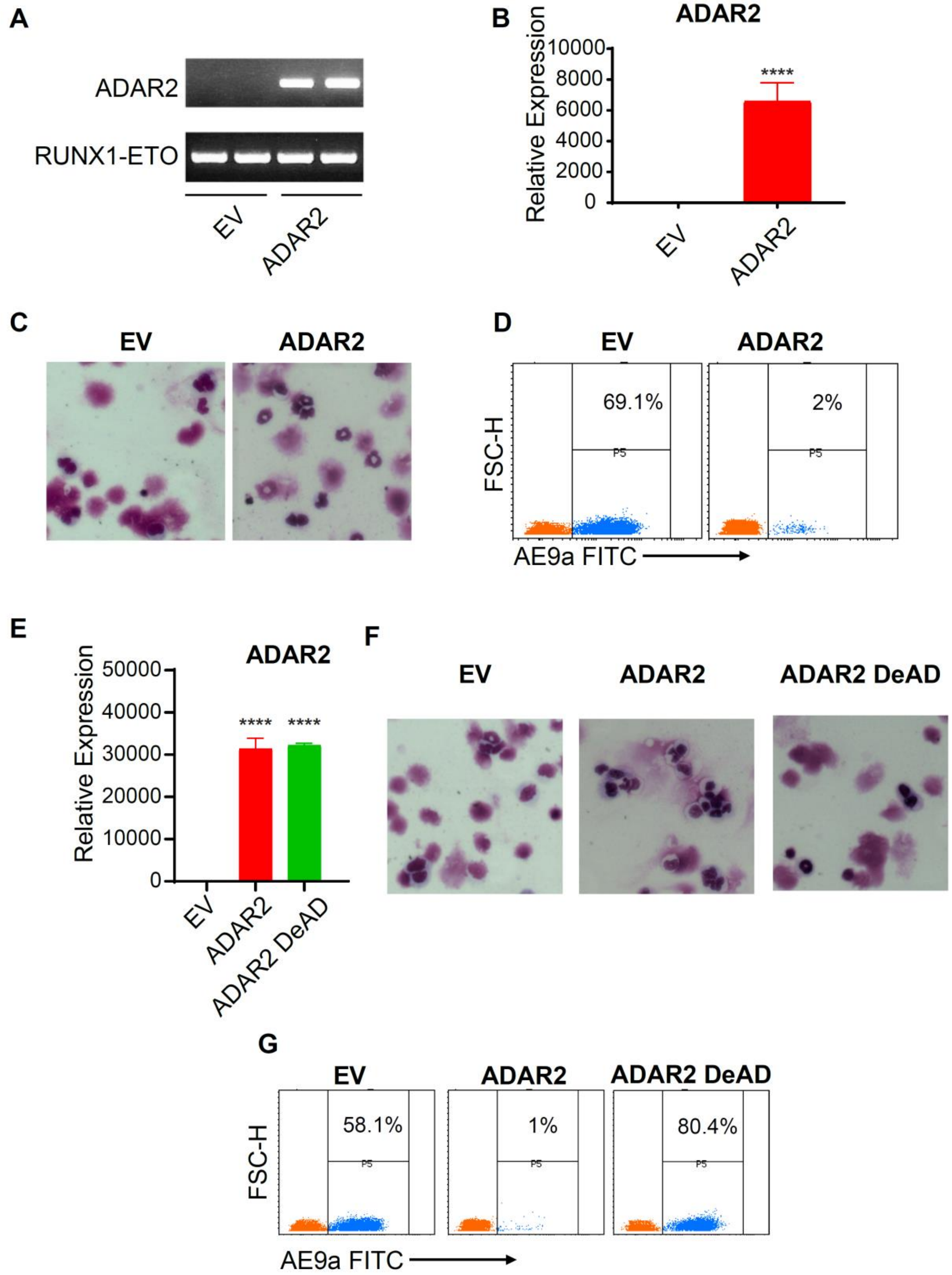

**Supplemental Figure 3. RNA editing capability of ADAR2 is essential for its repression of leukemogenesis in an AE9a mouse model**

**A, B)** RT-PCR A) and qPCR B) analyses of *ADAR2* transcript in the same samples as described in Fig. 4C. Data are presented as the mean ± SD of technical triplicates from a representative experiment in B). (****$p < 0.0001$, two-tailed Student's t-test.)

**C)** Representative pictures of Wright-Giemsa staining of bone marrow cells in the same recipients as described in Fig 4E.

**D)** Representative flow cytometric analyses of total bone marrow cells in the same recipients as described in Fig 4E.

**E)** qPCR analyses of *ADAR2* transcript in BM cells from the same recipients as described in Fig 4F. Data are presented as the mean ± SD of technical triplicates from a representative experiment of three independent experiments. (****$p < 0.0001$, two-tailed Student's t-test.)

**F)** Representative pictures of Wright-Giemsa staining of bone marrow cells in the same recipients as described in Fig 4H.

**G)** Representative flow cytometric analyses of total bone marrow cells of total bone marrow cells in the same recipients as described in Fig 4H.
